# Experimental warming reduces the diversity and functional potential of the *Sphagnum* microbiome

**DOI:** 10.1101/194761

**Authors:** Alyssa A. Carrell, Max Kolton, Melissa J. Warren, Dale A. Pelletier, Jennifer B. Glass, Joel E. Kostka, Paul J. Hanson, David J. Weston

**Affiliations:** Bredesen Center for Interdisciplinary Research and Graduate Education, University of Tennessee, Knoxville, TN 37996, USA; Biosciences Division, Oak Ridge National Laboratory, Oak Ridge, TN, 37831, USA; School of Biology, Georgia Institute of Technology, Atlanta, GA 30332, USA; School of Earth and Atmospheric Sciences, Georgia Institute of Technology, Atlanta, Georgia, 30332, USA; Environmental Sciences Division, Oak Ridge National Laboratory, Oak Ridge, TN, 37831, USA; Climate Change Science Institute, Oak Ridge National Laboratory, Oak Ridge, TN, 37831, USA

**Keywords:** *Sphagnum* microbiome, warming experiment, diazotroph diversity, simulated climate change, moss, microbial community

## Abstract

Climate change may reduce biodiversity leading to a reduction in ecosystem productivity. Despite numerous reports of a strong correlation of microbial diversity and ecosystem productivity, little is known about the warming effects on plant associated microbes. Here we explore the impact of experimental warming on the microbial and nitrogen-fixing (diazotroph) community associated with the widespread and ecologically relevant *Sphagnum* genus in a field warming experiment. To quantify changes in the abundance, diversity, and community composition of *Sphagnum* microbiomes with warming we utilized qPCR and Illumina sequencing of the 16S SSU rRNA and *nifH* gene. Microbial and diazotroph community richness and Shannon diversity decreased with warming (p<0.05). The diazotroph communities shifted from diverse communities to domination by primarily *Nostocaceae* (25% in control samples to 99% in elevated temperature samples). In addition, the nitrogen fixation activity measured with the acetylene reduction assay (ARA) decreased with warming treatment. This suggests the negative correlation of temperature and microbial diversity corresponds to a reduction in functional potential within the diazotroph community. The results indicate that climate warming may alter the community structure and function in peat moss microbiomes, with implications for impacts to host fitness and ecosystem productivity, and carbon uptake potential of peatlands.

## Introduction

Climate change represents a large threat to the function and stability of ecosystems, potentially leading to altered abundance range shifts (Parmesan, 2006), and species extinction (Parmesan, 2006; Bestion *et al.*, 2015) that ultimately result in decreased biodiversity. Despite years of research on the importance of diversity in driving the productivity and function in numerous ecosystems (Tilman *et al.*, 2012; Liang *et al.*, 2016; Kolton *et al.*, 2017; Laforest-Lapointe *et al.*, 2017), the relationship of warming and biodiversity remains unclear in many ecosystems. The majority of research on biodiversity and warming has focused mainly on multicellular eukaryotic organisms with little attention to the prokaryotes associated with them, but recent work has highlighted the key role that microbial biodiversity may play in determining the ecological response of ecosystems to warming (Bardgett & Putten, 2014; Bestion *et al.*, 2017).

Plant-microbe symbioses are widespread and ecologically important host-microbe associations. Plant-associated microbiomes have direct roles in ecosystem functioning through effects on carbon (Lu *et al.*, 2006; Knief *et al.*, 2012) and nitrogen cycles (Vile *et al.*, 2014; Moyes *et al.*, 2016). Plant microbial communities are structured by biotic factors (Bragina *et al.*, 2012; Berg *et al.*, 2014; Edwards *et al.*, 2015) and abiotic factors (Bulgarelli *et al.*, 2012; Lundberg *et al.*, 2012; Carrell & Frank, 2014; Edwards *et al.*, 2015) and have been found to be susceptible to environmental perturbations such as drought (Santos-Medellín *et al.*, 2017), nitrogen deposition (Gschwendtner *et al.*, 2016), and salinity (Yang *et al.*, 2016). Plant microbiomes also affect host plant health and productivity (Berendsen *et al.*, 2012; Chaparro *et al.*, 2012; Berg *et al.*, 2014), with more productive and healthy plants supporting greater microbial diversity (van der Heijden *et al.*, 2008; Berendsen *et al.*, 2012; Bever *et al.*, 2013; Agler *et al.*, 2016; Delgado-Baquerizo *et al.*, 2016; Kolton *et al.*, 2017). Despite the importance of microbes to plant function and ecosystem processes, and the sensitivity of plant-microbial symbioses to environmental disturbances, the response of plant associated-microbial diversity to climate warming is not well understood.

*Sphagnum* mosses play a large role in the global carbon cycle and are considered to be particularly vulnerable to climate change (McGuire *et al.*, 2009; Turetsky *et al.*, 2012). These bryophytes are inhabited by diverse microbes (Opelt *et al.*, 2007; Kostka *et al.*, 2016) with direct roles in the carbon cycle through methane oxidation (Raghoebarsing *et al.*, 2005; Kip *et al.*, 2010; Bragina *et al.*, 2013a), as well as other important ecosystem functions (Kostka *et al.*, 2016) such as nitrogen fixation (Bragina *et al.*, 2011, 2013b, 2014; Vile *et al.*, 2014; Warren *et al.*, 2017)(Bragina *et al.*, 2011, 2013a, 2014; Vile *et al.*, 2014; Warren *et al.*, 2017) that enables plant growth under nitrogen-limited conditions characteristic of the bogs where these mosses are found. Warming experiments have demonstrated that elevated temperature causes a reduction of *Sphagnum* biomass (Turetsky *et al.*, 2012). Moreover a recent study demonstrated elevated temperature may have both negative and positive impacts on *Sphagnum* microbial functional groups, which may destabilize carbon cycling in peatlands (Jassey *et al.*, 2013), but the effect of temperature on the community composition and diversity of Sphagnum microbiomes remains unknown.

In this study, we investigated the impact of experimental warming on the microbial community associated with *Sphagnum.* The objective of this study was to quantify changes in the abundance, diversity, and community composition of *Sphagnum* microbiomes with increased temperatures in the Spruce and Peatland Responses Under Changing Environments (SPRUCE) experiment (Hanson *et al.*, 2017) which provided in situ field warming treatments from ambient to +9°C at the S1-Bog of the Marcell Experimental Forest in northern Minnesota (Kolka et *al.*, 2011). The study focused on the nitrogen-fixing (diazotroph) functional guild that enables plant growth under the extreme nutrient-limited conditions characteristic of ombrotrophic bog ecosystems (Limpens & Heijmans, 2008; Larmola *et al.*, 2014; Vile *et al.*, 2014).

## Materials and Methods

### Experimental site and warming experiment

The SPRUCE experiment at the S1 bog on the Marcell Experimental Forest (Hanson *et al.*, 2017) employs a whole-ecosystem warming approach to produce nominal warming treatments of +0, +2.25, +4.5, +6.75 and +9 °C for a *Picea mariana* – *Sphagnum* spp. raised bog ecosystem. The experiment includes ten 12-m diameter plots with open-top enclosures (enclosed) and two ambient 12-m diameter plots without enclosures (non-enclosed). Briefly, the warming methodology combining air warming with deep-peat-heating from mild electrical resistance heaters to produce target warming levels superimposed over the natural diurnal and seasonal variability (Hanson et al. 2017). The experiment is located in the S1-Bog on the Marcell Experimental Forest (Kolka *et al*, 2011). The S1 Bog is an acidic and nutrient-deficient ombrotrophic *Sphagnum*-dominated peatland bog (surface pH≤4.0). The average means of annual precipitation and air temperature are 768 mm and 3.3°C respectively (Sebestyen *et al.*, 2011).

#### Sampling

To characterize the *Sphagnum* microbiome responses to warming, individual *Sphagnum* stems were randomly collected within each plot in June 2016 following continuous whole-ecosystem warming initiated in August of 2015. Samples were overnight shipped on ice to Oak Ridge National laboratory. Upon arrival, a subset of samples was shipped on ice overnight to Georgia Institute of Technology for ARA and the remaining plants were immediately pulverized with sterile mortar and pestle in liquid nitrogen for DNA extraction.

### DNA extraction, PCR and DNA sequencing

To characterize the abundance and community composition of *Sphagnum* microbiomes, DNA was extracted from 50 mg of each pulverized *Sphagnum* sample using a MoBio PowerPlant Plant Kit (MoBio, Carlsbad, CA, USA). Extracted DNA was frozen and shipped on dry ice to Georgia Institute of Technology for amplification and sequencing.

The diversity and composition of *Sphagnum* associated microbial communities was determined by applying a high-throughput sequencing-based protocol that targets PCR-generated amplicons from the V4 variable regions of the 16S rRNA gene using the primer set 515F (5′-GTGCCAGCMGCCGCGGTAA-3′) and 806R (5′-GGACTACHVGGGTWTCTAAT-3′) as previously described (Wilson *et al.*, 2016; Kolton *et al.*, 2017). The diversity and composition of diazotrophic communities were assessed by targeting *nifH* (encoding the nitrogenase reductase subunit) as a molecular marker for nitrogen-fixing microorganisms. Primers IGK3 (5’-GCIWTHTAYGGIAARGGIGGIATHGGIAA-3’) and DVV (5’-TIGCRAAICCICCRCAIACIACRTC-3’) were employed to generate 396 bp PCR products (Gaby & Buckley, 2014). The 16A SSU rRNA and *nifH* amplicons were barcoded with unique 10-base barcodes (Fluidigm Corporation), and sequenced on an Illumina MiSeq2000 platform at the Georgia Institute of Technology following standard protocols (Caporaso *et al.*, 2012; http://www.earthmicrobiome.org/emp-standard-protocols/16s/; Gilbert *et al.*, 2010; Gaby *et al.*, 2017, submitted).

### Sequence processing and analysis

First, Illumina-generated 16S SSU rRNA and *nifH* gene amplicon sequences were paired with PEAR (Zhang *et al.*, 2014) and primers were trimmed with the software Mothur v1.36.1 (Schloss *et al.*, 2009). Resulting sequences were quality filtered using a Phred quality score Q30 and Q25 for 16S SSU and nifH respectively using the standard QIIME 1.9.1 pipeline (Caporaso *et al.*, 2010). Sequences were clustered into operational taxonomic units (OTUs) by using UCLUST algorithm with a threshold of 97% identity. Representative sequences were aligned using PyNAST (Caporaso *et al.*, 2010) against the Greengenes core set for 16S SSU and against *nifH* gene alingment (DeSantis *et al.*, 2006; Gaby & Buckley, 2014). Taxonomies of these high-quality sequences were annotated to the Greengenes database (release 13_8) (DeSantis *et al.*, 2006) or a manually curated *nifH* database (Gaby & Buckley, 2014) using the RDP classifier (Wang *et al.*, 2007) with a minimum confidence threshold of 50%. The 16S SSU rRNA sequences classified as “chloroplast” or “mitochondria” were removed from the alignment. An approximately maximum-likelihood tree was constructed from the aligned of bacterial representative sequences, using FastTree (Price *et al.*, 2009). Prior to conducting diversity analyses, OTUs were rarefied to 3500 reads per sample for 16S SSU rRNA amplicons and 1500 reads per sample for *nifH* amplicons. The OTU-based alpha diversity was calculated based on the total number of phylotype (observed richness) and on Shannon’s diversity index (H′). Faith’s phylogenetic diversity (PD) was calculated to assess phylogenetic based alpha diversity. The OTU-based beta diversity indices were estimated based on Bray–Curtis distances.

The Illumina-generated 16S SSU rRNA and *nifH* gene amplicon sequences have been deposited in the BioProject database, (ncbi.nlm.nih.gov/bioproject) under accession PRJNA407792 and PRJNA407800 respectively.

### Quantitative PCR amplification

All quantitative polymerase chain reaction assays were performed in triplicates using the StepOnePlus platform (Applied Biosystems, USA) and PowerUp SYBR Green Master Mix (Applied Biosystems, USA). Absolute quantification of 16S SSU rRNA and *nifH* genes were conducted with primer pairs 331F (5’-CCTACGGGAGGCAGCAGT-3’)/518R (5’-ATTACCGCGGCTGCTG-3’) and PolF (5’-TGCGAYCCSAARGCBGACTC-3’) /PolR (5’-ATSGCCATCATYTCRCCGGA-3’) respectively (Muyzer & Waal, 1993; Poly *et al.*, 2001). The 16S SSU rRNA quantification reaction was carried out in 20 μl containing 7.8 μl of PCR grade water, 0.1 μl of each primer (final concentration 0.5 μM), 10 μl of PowerUp SYBR Green Master Mix (Applied Biosystems, USA) and 2 μl of sample DNA. The cycling program included 2 min at 50 °C, 2 min at 95 °C, followed by 40 cycles of 95 °C for 15 s, 55 °C for 15 s and 72 °C for 1 min. The *nifH* gene quantification reaction was carried out in 20 μl containing 6.8 μl of PCR grade water, 0.6 μl of each primer (final concentration 0.3 μM), 10 μl of PowerUp SYBR Green Master Mix (Applied Biosystems, USA) and 2 μl of sample DNA. The cycling program included 2 min at 50 °C, 2 min at 95 °C, followed by 45 cycles of 95 °C for 15 s, 63 °C for 1 min. Amplification specificity was studied by melting curve analysis of the PCR products, performed by ramping the temperature to 95 °C for 15 s and back to 60 °C for 1 min, followed by increases of 0.15 °C s^-1^ up to 95 °C. Melting curve calculation and determination of Tm values were performed using the polynomial algorithm function of StepOnePlus Software (Applied Biosystems, USA). In all experiments, negative controls containing no template DNA were subjected to the same qPCR procedure to exclude or detect any possible DNA contamination. Standard curves were obtained with serial dilution of standard plasmids containing target *Escherichia coli* k12 16S rRNA or *Azotobacter vinelandii nifH* gene fragments as the insert. The abundance of standard plasmid inserts ranged from 2.97 × 10^3^ to 2.97 × 10^9^ (bacterial 16S SSU rRNA gene) or 24.2 to 2.42 × 10^6^ (*nifH* gene).

### Acetylene reduction assay

To determine the effect of warming on nitrogen fixation activity, fresh tissue from the ambient and warming plots exposed to the highest temperatures (+9°C) were interrogated using the acetylene reduction assay (ARA) as previously described (Warren *et al.*, 2017). Briefly, samples of *Sphagnum* were collected from ambient enclosed and non-enclosed plots and +9°C enclosed plots in triplicate and stored at 4°C until the start of incubations. A 1.0-1.5 g sample of green-only *Sphagnum* was placed into 35 ml glass serum bottles, stoppered with black butyl stoppers, sealed with an aluminum crimp seal, and 10% headspace was replaced with 10% room air or with 10% C_2_H_2_. Controls that were not amended with C_2_H_2_ did not produce detectable ethylene. All treatments were incubated for one week in the light at 25°C. A gas chromatograph with flame ionization detector (DRI Instruments, Torrance, CA, USA) equipped with a HayeSep N column was used to quantify ethylene (C_2_H_4_). The accumulation of C_2_H_4_ was determined twice daily until C_2_H_4_ production was linear (∼3 days). Samples were dried at the end of incubations at 80°C for 48 hours to determine dry weight for normalization of ARA rates.

### Data analysis

Statistical analysis was conducted in R (R Core Team, 2015). Warming effects on microbiome community composition were assessed with a Spearman *Rho* test between warming treatments and a heatmap was generated from the relative abundance of distinct OTUs that showed significant differences (p<0.05) and had >0.1% relative abundance in at least a single treatment. General Linear Models (GLMs) were used to evaluate the effects of warming on microbial diversity measurements of enclosed plots. A Mann-Whitney test was used to compare diversity between ambient plots, with or without an enclosure structure. Beta diversity was visualized using non-metric multidimensional scaling ordination (NMDS) from Bray-Curtis similarity distances. Analysis of similarities (ANOSIM) and permutational multivariate analysis of variance (PERMANOVA), each with 999 permutations, were used to determine if beta diversity differed significantly among treatments.

## Results

### Response of microbiome abundance, community composition, and diversity to warming

The overall microbial abundance as determined by qPCR did not vary by warming treatment (*p*=0.2; Table 1).

**Table 1.** Effect of warming on bacterial and diazotroph gene abundances. Triplicate samples from duplicate plots of each warming treatment were used to calculate the average absolute abundance with standard error of bacterial (16S SSU rRNA) and diazotroph (*nifH*) gene abundance of *Sphagnum* bacteria.

| Assay | +0°C | +2.25°C | +4.5°C | +6.75°C | +9°C |
| --- | --- | --- | --- | --- | --- |
| Bacterial 16S SSU rRNA gene abundance<br>(10<sup>8</sup> per g *Sphagnum* tissue) | 15.18 (3.86) | 14.58 (2.79) | 9.56 (2.71) | 8.83 (4.08) | 9.37 (1.55) |
| Diazotroph *nifH* rRNA gene abundance<br>(10<sup>8</sup> per g *Sphagnum* tissue) | 0.05 (0.03) | 0.03 (0.004) | 0.03 (0.003) | 0.03 (0.008) | 0.02 (0.002) |

The *Sphagnum* microbiome communities were dominated by *Proteobacteria* (62%) and *Acidobacteria* (17%), with smaller contributions from candidate division WPS-2 (4%), *Cyanobacteria* (4%), *Bacterioidetes*, (3%) *Verrucomicrobia* (2%), and *Actinobacteria* (1%) with Cyanobacteria varying across warming treatments though not significant (Fig. 1). The *Proteobacteria* were dominated by the order *Rhodospirillales* (33%) followed by *Caulobaceterales* (7%), *Xanthomondales* (8%), and *Burkholderiales* (3%). Despite the dominance in major phyla and genera groups across treatment, several OTUs varied significantly across warming treatment (Table S1). Cyanobacteria in the *Nostocaceae* family, OTU 278041 most similar to *Nostoc sp.* and OTU 4242238 most similar to *Cylindrospermum sp.*, increased in relative abundance from 0.4 to 4.1% and from 0 to 1%, respectively, across all warming treatments (*p*=0.04). Warming treatments had a varied effect on *Acetobacteraceae* with relative abundance decreasing in +2.25°C and +4.5°C treatments but returning to similar abundances in +6.75 and +9°C treatments.

**Figure 1.**
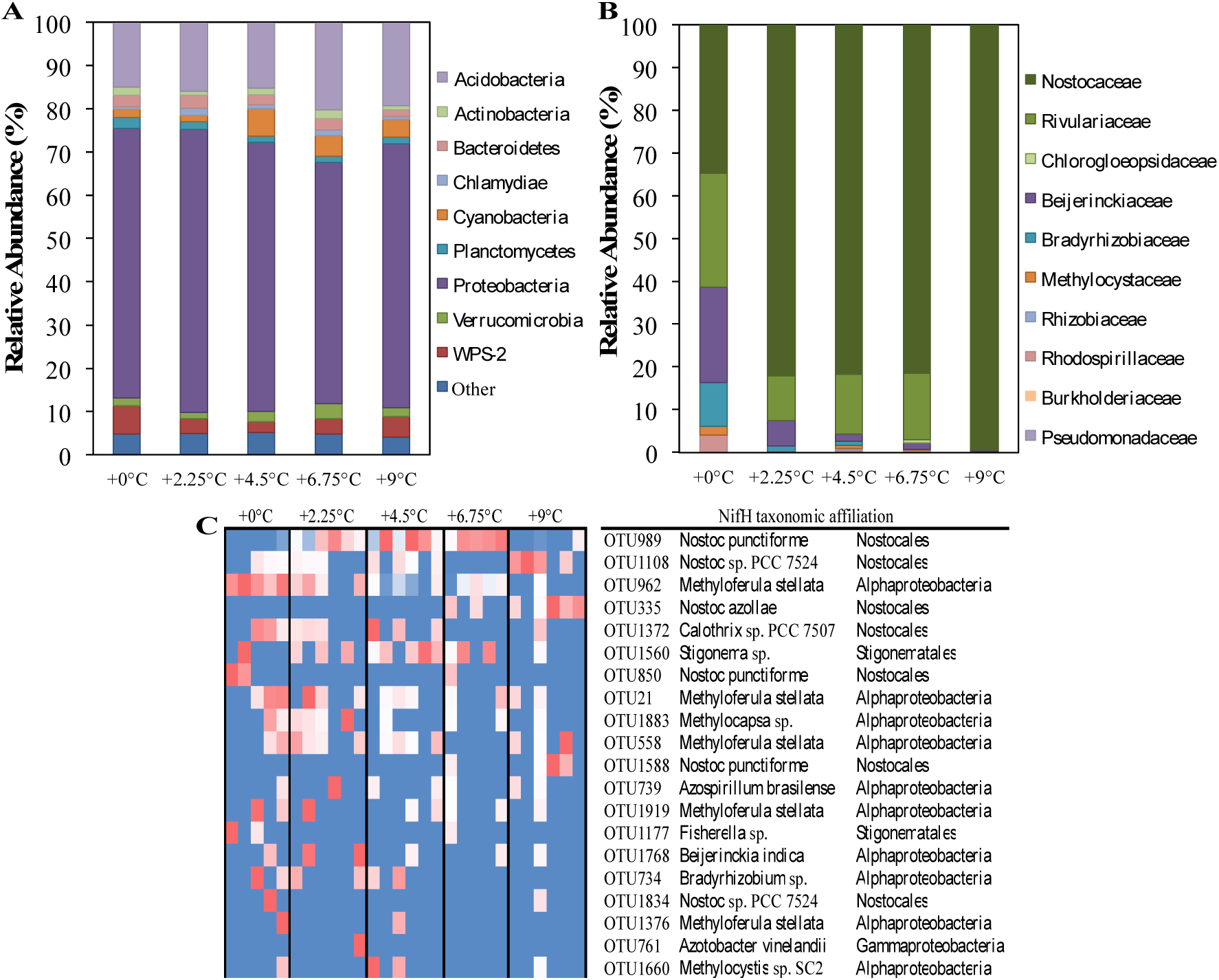
Effect of warming on overall microbial and diazotroph community composition in *Sphagnum* microbiomes. Relative abundance of 16S SSU rRNA or nifH gene sequences was determined at various taxonomic levels from triplicate samples collected in duplicate enclosures for each treatment plot. Average relative abundance of 16S SSU rRNA gene amplicons (A) at the phylum level and nifH gene amplicons (B) at the family level from each warming treatment. A heatmap was generated of top the 20 *nifH* phylotpes with BLAST taxonomic family and species identity (C). For each OTU, the highest abundance is indicated by dark red, intermediate is white, and lowest abundance is blue with a color gradient for the remaining values.

The richness and phylogenetic diversity of *Sphagnum* microbiomes decreased with warming. Observed richness and Shannon index decreased with warming (*p*<0.05), while phylogenetic diversity decreased with warming treatment but was only significant at *p*=0.08 (Fig. 2, Table 2). *Sphagnum* bacterial communities were structured by warming treatments (*p<*0.003) with Bray-Curtis distance similarity higher within treatment than between treatments (Fig. S1). Percent similarity for all samples was 52% (standard deviation = 5%) with a range of 31-65% similarity. (R^2^=0.3, *p*=0.004).

**Figure 2.**
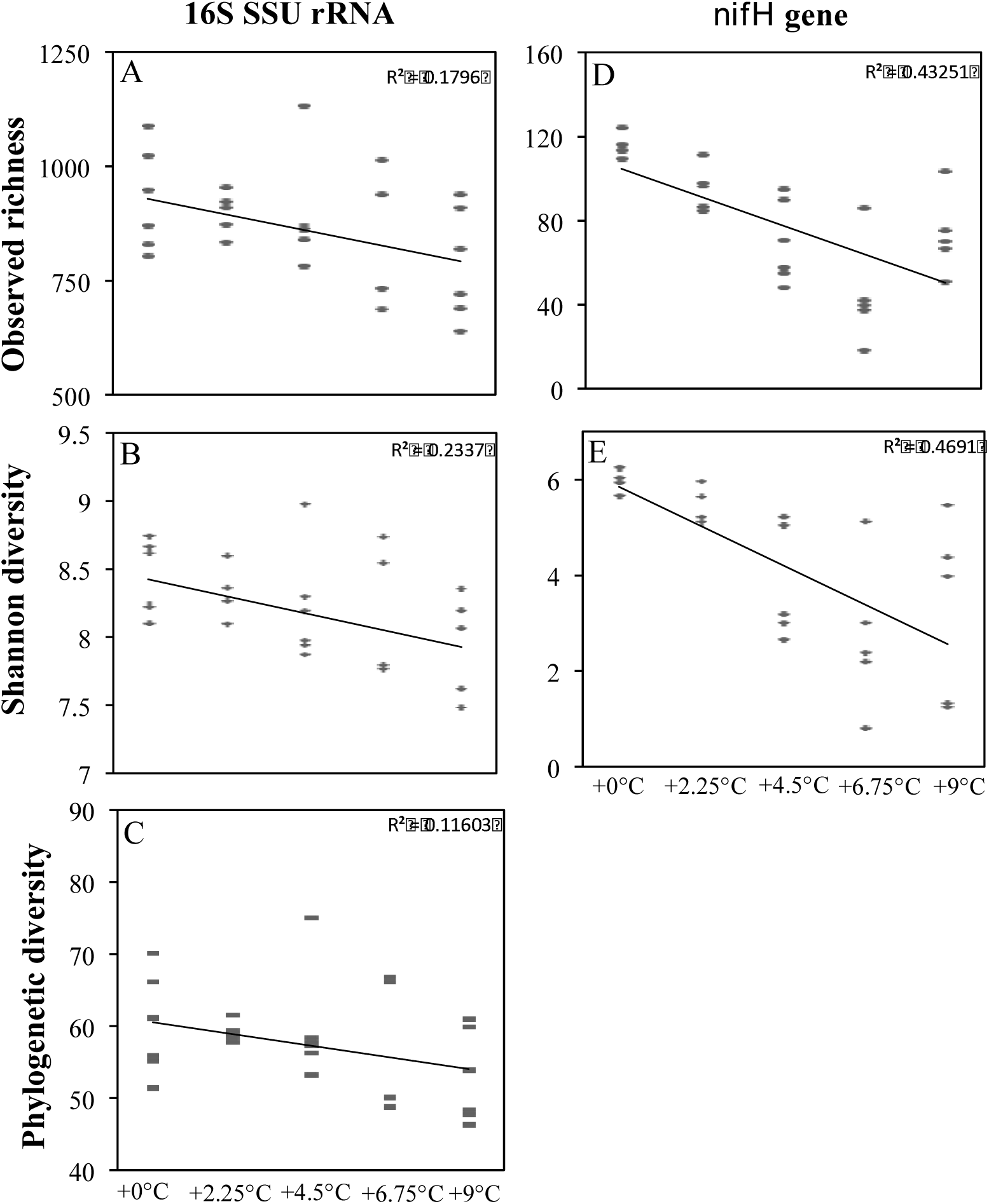
Effect of warming on alpha diversity of *Sphagnum* bacterial and diazotroph communities. Triplicate samples collected in duplicate enclosures for each treatment plot were used.to calculate observed Operational Taxonomic Units (OTUs) (A and D), Shannon’s diversity (B and F), and phylogenetic diversity (C) of 16S SSU rRNA gene (A-C) sequences rarefied to 3500 sequences per sample and *nifH* gene (D and F) sequences rarefied to 1500 sequences per sample.

**Table 2.**
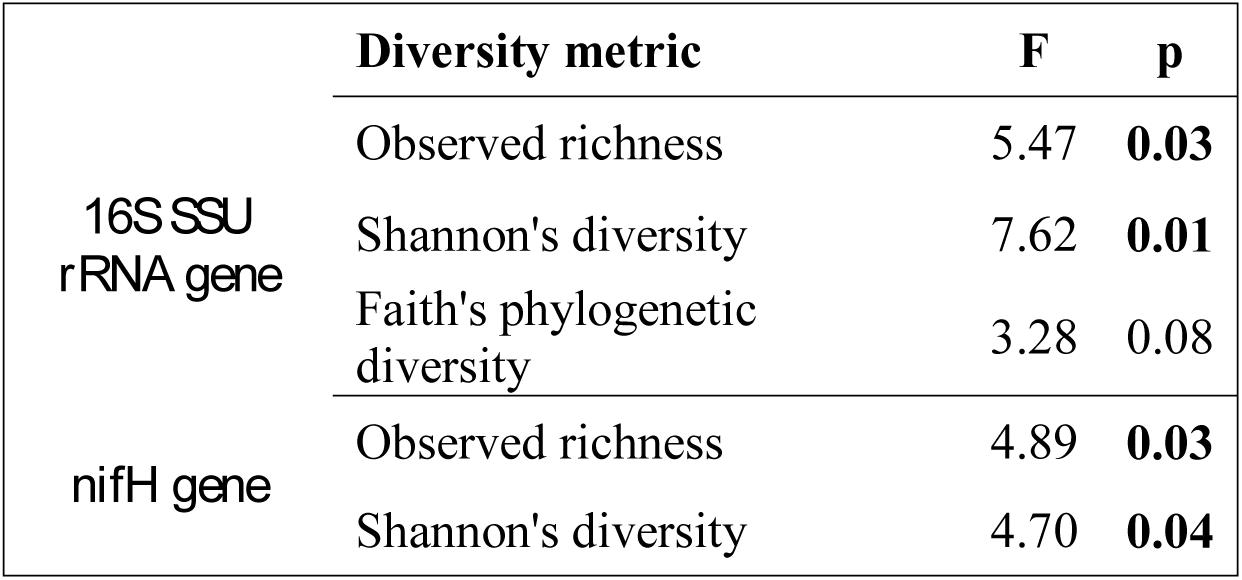
Effect of warming on the alpha diversity of *Sphagnum* bacteria. General Linear Models (GLMs) were used to evaluate the effects of warming on microbial diversity measurements of enclosed plots. Triplicate samples from duplicate plots of each warming treatment were used to measure observed OTU richness, Shannon’s diversity, and Faith’s phylogenetic diversity of *Sphagnum* bacterial 16S SSU rRNA genes across warming enclosure treatment plots. Significance metrics are indicated in bold (p<0.05).

### Response of diazotroph abundance, diversity, community composition and function to warming

The abundance of diazotrophs as determined by qPCR of *nifH* genes significantly decreased (*p*=0.004) with increasing temperature (Table 1). All *nifH* gene profiles were dominated by the phyla *Cyanobacteria* (60-100%) and *Proteobacteria* (0.5-40%) with *Cyanobacteria* increasing in abundance and *Proteobacteria* decreasing with warming treatments. Abundant members of the *Cyanobacteria* phylum were comprised of *Nostocaceae* (25-99%), *Rivulariaceae* (0-27%), and *Chlorogloeopsdidaceae* (0-0.7%), with *Nostocaceae* becoming more dominant with warming (Fig. 1). The *Rhizobiales* (0.1-35%) and *Rhodospirillales* (0-4%) were detected in abundance from the *Proteobacteria* phylum, with relative abundance decreasing across warming treatments. To provide greater resolution into shifts in diazotroph populations, an OTU heatmap was generated from the top 20 OTUs of each treatment (Fig. 1). Notably, ambient warming plots were largely dominated by sequences most similar to the genera *Methyloferula* (17-40%) and *Calothrix* (0-32%) which both decreased across warming treatments: +2.25°C (0-25%), +4.5°C (0-7%), +6.75°C (0-6%), and +9°C (0-3%). With increased warming, sequences closely related to the genus *Nostoc* became more dominant though different *Nostoc* species dominated across each temperature treatment. Sequences most similar to *Nostoc punctiforme* dominated the +2.25°C (20-80%), +4.5°C (26-88%), and +6.75°C (46-83%) treatments while +9°C was dominated by *Nostoc* sp. PCC7524 (0-100%).

Warming reduced the richness and diversity of the diazotroph community (p<0.05, Table 2), although each treatment did not respond equally. When compared to +0°C, diazotroph richness at +2.25°C and + 4.5°C decreased by 30% and 54%, respectively, while richness in the +6.75°C and +9°C plots only decreased by 14% richness (Fig. 2, Table S2). Shannon indices followed a similar pattern with a reduction in diversity of 27% in the +2.25°C plots, 52% in the + 4.5°C plots, 18% in the 6.75°C plots and only 3% in the +9°C plots (Fig. 2). The diazotroph community was structured by temperature treatment in that samples from the same treatment clustered closer to one another than other treatments (R^2^=0.3546, *p*=0.041). However, the clustering was not incremental with diazotroph communities from 0°C and 9°C clustering closer to one another than with 6.75°C (Fig. S1).

Nitrogen fixation rates determined by ARA showed considerable variability within warming treatments, with some samples showing no detectable activity while others had rates as high as 172 nmol g^-1^ hr^-1^. Average rates of nitrogen fixation decreased by ∼50% from +0°C (47 ±9 nmol g^-1^ hr^-1^) to +9°C (21 ± 6 nmol g^-1^ hr^-1^), but the decline was only significant at *p*=0.1, due to variation between replicates (Fig. 3).

**Figure 3.**
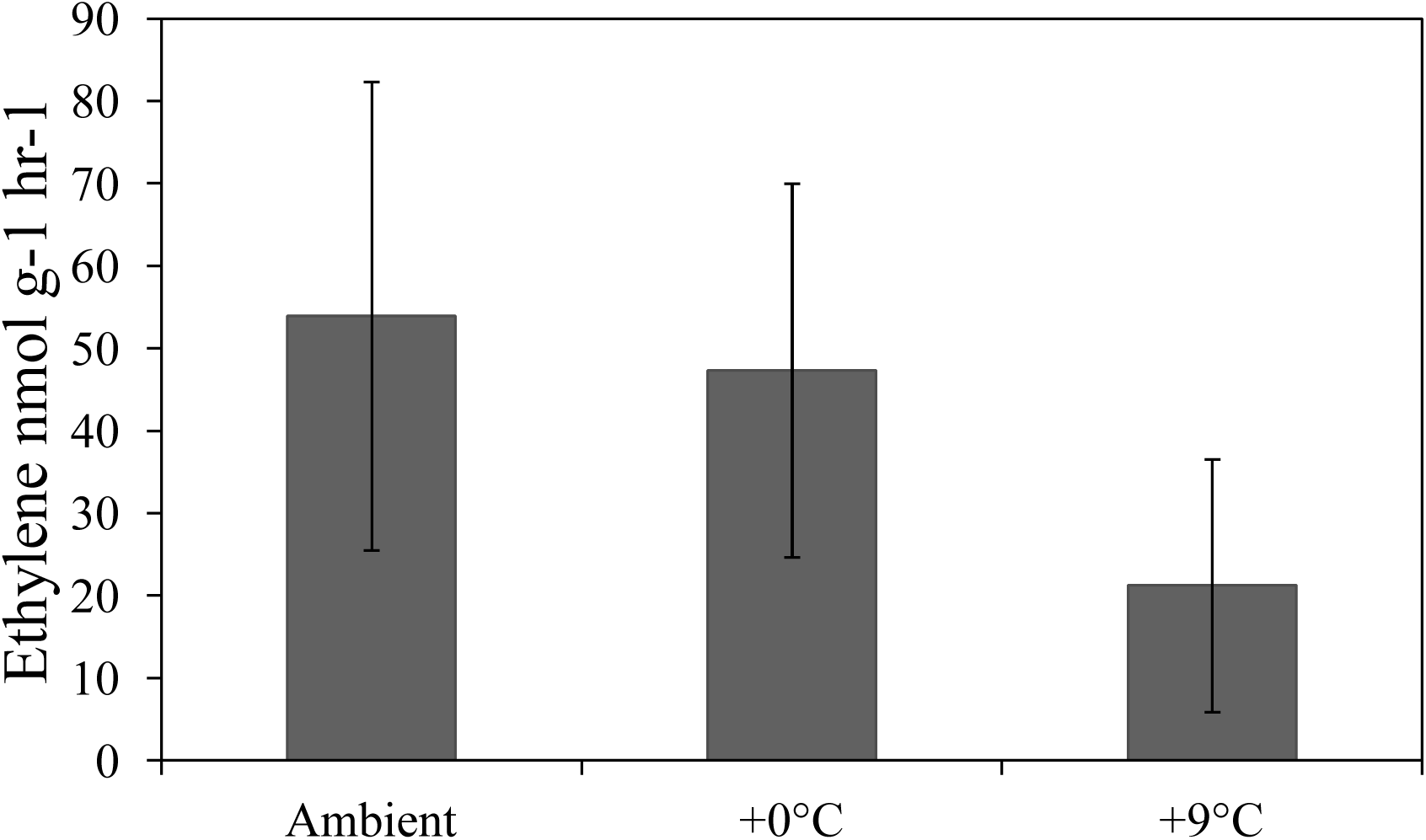
Effect of warming on nitrogenase activity. Triplicate samples from duplicate plots of enclosed ambient (0°C) and +9°C and non-enclosed ambient (ambient) treatments were used to measure potential nitrogenase activity with the acetylene reduction assay. Error bars represent 1 standard deviation.

### Experimental enclosure affect

To test if the presence of the experimental structure had a significant impact on *Sphagnum* general bacterial and diazotroph community composition and diversity, we measured 16S SSU rRNA and *nifH* genes of *Sphagnum* in ambient plots without an enclosure (ambient) and ambient plots with an enclosure warmed at +0°C above outside ambient conditions. We found that the enclosure had no statistical effect on 16S SSU rRNA and *nifH* gene composition, abundance, diversity, richness or evenness (Figure S2, Tables S1, S2). Temperature did not significantly change community structure for either 16S SSU rRNA (R^2^=0.02, *p*=0.4) or *nifH* genes (R^2^=0.01, *p*=0.6).

## Discussion

Determining the potential effects of climate drivers such as temperature on *Sphagnum* microbiomes is an important step toward effectively predicting the response of ecosystem function in ombrotrophic bogs to climate change. Here we demonstrate that temperature strongly influences general microbial and diazotroph community structure and diversity. Additionally, *Sphagnum* microbiome communities from ambient plots without enclosure were not significantly different in microbiome or diazotroph composition, abundance, or diversity, than *Sphagnum* microbiome communities in plots with enclosures, indicating that differences between temperature treatments were not an artifact of the experimental warming structure.

### Warming effects on overall microbiome communities

The *Proteobacteria*, *Acidobacteria*, and *Cyanobacteria* dominated all samples, and have been found to dominate *Sphagnum* in other bog systems (Bragina et al., 2014). Despite consistent dominance by the same phyla, overall community structure differed by warming treatments, likely due to variation at a lower taxonomic level. We did see variation in species within bacterial families, possibly as a result of differential temperature optima of bacterial species. Overall, observed richness, diversity and phylogenetic diversity were negatively correlated with temperature. Phylogenetic diversity is a divergence based method that has been described as more powerful than qualitative measurements given the correlation of 16S SSU rRNA similarity and phenotypic similarities in microbial key features such as metabolic capabilities or other functions (Lozupone & Knight, 2008). This would suggest that while we see a reduction in overall phylotype counts, we also see a reduction in metabolic capabilities.

A reduction of microbial diversity may make ecosystems more susceptible to environmental perturbations and when considering additional perturbations such as N deposition or different precipitation patterns, these communities may be even more impacted (Aanderud *et al.*, 2013). Here we found a reduction of richness and diversity in both the general microbial community and diazotroph community. Indeed a reduction of richness and evenness of microbial communities in other ecosystems such as soil or rhizosphere, were associated with a decrease in ecosystem functioning such as nutrient cycling (Philippot *et al.*, 2013; Wagg *et al.*, 2014), plant productivity (Bell *et al.*, 2005; van der Heijden *et al.*, 2008; Lau & Lennon, 2011; Fierer *et al.*, 2013) and plant resilience against pathogen invasion (Jousset *et al.*, 2011; Mendes *et al.*, 2011). Moreover, reduction in microbial diversity is frequently associated with reduced activation of plant defense systems (Mendes *et al.*, 2011, 2013; Berendsen *et al.*, 2012). Additionally, *Sphagnum* mosses have been found to harbor potential latent plant pathogens and in many organisms disease outbreaks are dependent on the abundance of pathogens and the diversity of microbiomes (Bragina *et al.*, 2011; Elad & Pertot, 2014; Tout *et al.*, 2015). Alternatively, a reduction in diversity could correspond to a loss of pathogenic taxa, which might be beneficial to host plants. Therefore, further study will be needed to determine the specific ecosystem functions that are mediated by the *Sphagnum* microbiome and impacted by warming.

### Warming effects on diazotroph communities

Nitrogen is essential to the growth and maintenance of *Sphagnum* plants and previous research revealed highly specific and diverse diazotrophs (Bragina *et al.*, 2013a)(Bragina *et al.*, 2013a) are a major source of N in *Sphagnum*-dominated peatlands (Lindo *et al.*, 2013; Larmola *et al.*, 2014; Vile *et al.*, 2014; Novak *et al.*, 2016). In corroboration of patterns in overall microbiome communities, diazotroph diversity and abundance were negatively correlated with temperature. This suggests that the reduction of microbial diversity may lead to a reduction of functional potential within the diazotroph functional guild. Within the diazotroph community, we found a shift in community composition with elevated temperature leading to a community dominated by primarily by *Nostoc* and void of diazotrophic methanotrophs. In addition, another filamentous cyanobacterium, *Stigonema,* was shown to decrease in relative abundance across temperature treatment to below detection in the +9°C treatment. Interestingly, *Nostoc* has been described as “cheaters” in the feather moss microbiome as it dominated the cyanobacterial community but had low *nifH* gene expression and thus not providing much nitrogen to the host. Conversely, *Stigonema* made up less than 29% of the cyanobacterial community but accounted for the majority of *nifH* gene expression suggesting *Stigonema* is responsible for the majority of fixed nitrogen (Warshan *et al.*, 2016). Though it is possible an observed reduction in nitrogen fixation may be attributed to the increase in the presence of a “cheater” and/or disruption of supportive metabolic pathways it cannot be concluded from our data that *Nostoc* is a cheater in our system. Concurrent with an increase in *Nostoc* relative abundance we found a decrease in diazotroph absolute abundance indicating that *Nostoc* may not be increasing in abundance but rather other microbial populations, such as the methanotrophs, are dropping out of the community.

### Diazotroph function

Nitrogen fixation activity and temperature were negatively correlated which may be due to plant-specific tolerance to water stress and desiccation given that nitrogen fixation associated with moss is influenced by moisture (Zielke *et al.*, 2002; Sorensen *et al.*, 2006; Sorensen & Michelsen, 2011). Additionally, oxygen level, photosynthetic activity (Warren *et al.*, 2017), and phosphorous (Rousk *et al.*, 2017) or nitrogen availability (Kox *et al.*, 2016) have also been found to limit diazotrophy in *Sphagnum* (Warren *et al.*, 2017). However, we observed a reduction in diazotroph absolute abundance indicating diazotrophs were not inactive but rather undetectable with our methods in elevated temperature treatments. Alternatively, this may be attributed to the diazotroph optimal temperature for nitrogen fixation (Gundale *et al.*, 2012) or a disruption in microbiome composition. The nitrogenase enzyme commonly contains molybdenum (Rousk *et al.*, 2017; Warren *et al.*, 2017) as its cofactor but may contain vanadium or iron in its place (Miller & Eady, 1988). Thus the change across temperatures could be attributed to altered metal availability. With a reduction in nitrogen fixation, *Sphagnum* may become more reliant on nitrogen provided by non-associative diazotrophs such as bacteria in the pore water or below peat. However, if the *Sphagnum* associated microbes are susceptible to elevated temperature, diazotrophs in the water may be even more so. Additionally, *Sphagnum* competition for other sources of nitrogen may disrupt free-living microbial communities causing larger consequences at the ecosystem level.

Though we found a general pattern of a reduction in potential rates of nitrogen fixation, it is important to note that acetylene inhibits the enzyme methane monooxygenase and thus the diazotrophy of methanotrophs. A recent study calibrated ARA with 15N incorporation and found a conversion factor of 3.9 for 15N_2_-to-ARA in the same bog as our experiment, indicating the presence of diazotrophic methanotrophs that were inhibited by acetylene (Warren *et al.*, 2017). In our study, the use of the conversion factor is inappropriate given the demonstration of an altered diazotroph community. While it is possible we have underestimated diazotroph activity, our observations of decreased nitrogen fixation activity with warming are supported by a decline in diazotroph abundance and the relative abundance of diazotrophic methanotrophs.

With warming induced reduction of diazotroph abundance and function, one might logically expect a decline in peatland ecosystems carbon storage capacity. The considerable accumulation of C as peat results from a long-term excess of Net Primary Productivity (NPP) of plants over peat decomposition. In peatlands a simple mass balance demonstrates N-deposition alone does not account for the N needed to support the observed NPP (Wieder *et al.*, 2010). A recent study demonstrated diazotrophs may account for 12-25 times more N than from atmospheric inputs alone, accenting the important link between diazotrophy and NPP (Vile *et al.*, 2014). *Sphagnum* has demonstrated differential NPP response to warming (Aerts *et al.*, 2006) but no studies have examined the *Sphagnum* microbial community and diazotroph responses to warming. Here we present data that suggests warming may disrupt the diazotroph community and function, which ultimately may reduce NPP or the accumulation of peat and therefore may be an important component to include in future *Sphagnum* and peatland response studies.

Microbial associates play an important role in *Sphagnum* health and growth as well as bog ecosystem functioning. In this study, we conducted a warming experiment to elucidate the temperature effects on *Sphagnum* microbiomes. We propose that climate warming may alter microbiome function as a result of decreased biodiversity. The consequences of decreased functional potential are not clear and merits future studies to determine how the alteration of overall microbiome and diazotroph function may scale to the ecosystem level. Such knowledge will provide a more comprehensive understanding of how climate may impact the future function of *Sphagnum* dominated bog ecosystems.

## Acknowledgements

The experiment were maintained as part of the SPRUCE project and supported by the U.S. Department of Energy’s Office of Science, Biological and Environmental Research (DOE BER). Oak Ridge National Laboratory is managed by UT-Battelle, LLC, for the U.S. Department of Energy under contract DE-AC05-00OR22725. Sample collection, processing and manuscript writing was supported by the Laboratory Directed Research and Development Program of Oak Ridge National Laboratory, managed by UT-Battelle, LLC, for the U. S. Department of Energy. Sequencing was supported by U.S. DOE BER under award numbers DE-SC0007144 and DE-SC0012088.

